# Threshold-crossing time statistics for gene expression in growing cells

**DOI:** 10.1101/2022.06.09.494908

**Authors:** César Nieto, Khem Raj Ghusinga, César Vargas-García, Abhyudai Singh

## Abstract

Many intracellular events are triggered by attaining critical concentrations of their corresponding regulatory proteins. How cells ensure precision in the timing of the protein accumulation is a fundamental problem, and contrasting predictions of different models can help us understand the mechanisms involved in such processes. Here, we formulate the timing of protein threshold-crossing as a first passage time (FPT) problem focusing on how the mean FPT and its fluctuations depend on the threshold protein concentration. First, we model the protein-crossing dynamics from the perspective of three classical models of gene expression that do not explicitly accounts for cell growth but consider the dilution as equivalent to degradation: (*birth-death process, discrete birth with continuous deterministic degradation*, and *Fokker-Planck approximation*). We compare the resulting FPT statistics with a fourth model proposed by us (*growing cell*) that comprises size-dependent expression in an exponentially growing cell. When proteins accumulate in growing cells, their concentration reaches a steady value. We observe that if dilution by cell growth is modeled as degradation, cells can reach concentrations higher than this steady-state level at a finite time. In the growing cell model, on the other hand, the FPT moments diverge if the threshold is higher than the steady-state level. This effect can be interpreted as a transition between noisy dynamics when cells are small to an almost deterministic behavior when cells grow enough. We finally study the mean FPT that optimizes the timing precision. The growing cell model predicts a higher optimal FPT and less variability than the classical models.

## I. Introduction

At the level of an individual cell, gene products exhibit stochastic fluctuations (noise) [1], [2]. This noise stems from the inherent randomness of the synthesis and degradation of specific gene products [3], [4] as well as from variability in cell-cycle dependent variables such as gene replication [5], cell size dynamics [6], molecule segregation at division [7], [8], and growth rate [9], [10]. Although stochastic models of gene-expression have been studied for quite some time, effects of global processes such as cellular growth, cell size regulation, etc. have been incorporated only recently to these models [11]–[14].

Both number and concentration of a gene-product are the main quantities of interest in gene-expression models and they are characterized by computing their probability distributions or a few lower-order statistical moments. Gene-product fluctuations are studied also by the first passage times (FPT). Formally, FPT is the time spent to reach a threshold being relevant in many cases [15]–[17]. For example, accumulation of a threshold number of molecules is thought to be a mechanism for cell division [18], [19] that can be modeled as an FPT problem [20]. Other kind of FPT problems such as bacterial sporulation [21], cell differentiation [22] and apoptosis [23] involve reaching a threshold concentration, not a molecule number.

The difference between reaching either an amount or a concentration is often not clear. Some studies [16], [24], [25], for instance, consider reaching concentration as equivalent to molecule accumulation since they assume that cell size does not change appreciably during the molecule accumulation [4], [26]. When cell size grows exponentially, FPT problems comprising concentration threshold are expected to have different properties from those comprising molecule number threshold.

To illustrate this effect, consider a cell whose size grows exponentially over time and production rate of a protein of interest scales with cell size [27]–[29]. In case of numbers threshold, the mean FPT is smaller for a cell with a larger size at birth [20]. On the contrary, for a concentration threshold, we expect a rather weak dependency between mean FPT and cell size at birth since both the production rate as well as the number of molecules required to achieve a set concentration scale with cell size. In other words, production is balanced by dilution of gene-product concentration in a growing cell. In most models, effects of deterministic dilution by growth are taken to be equivalent to a random process such as either degradation or segregation at division [30]– [33]. It is however unclear whether explicitly accounting for cell-size and growth would result in the same FPT statistics as the models that do not do it.

Here, we study FPT statistics (mean FPT and noise in FPT, quantified by the squared coefficient of variation) for the protein concentration threshold-crossing in exponentially growing cells with protein synthesis rate proportional to the size. We compare these FPT statistics with the statistics of three other models that do not consider cell size dynamics: the discrete birth-death Poisson process [34], discrete stochastic production and continuous dilution [30], and the continuous process with Brownian noise by the Fokker-Planck equation [32]. An important finding is that the model that explicitly accounts for cell size dependency on gene expression has lower noise in FPT than the other three models.

## II. Modelling the FPT distribution for thresholding

Let *c*(*t*) denote the concentration of a gene-product at time *t*. Assuming *c*(0) = 0, the first passage time *T* to cross a threshold *C* is given by

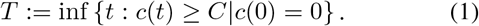

The probability distribution and moments of *T* depend upon the description of *c*(*t*). Below, we describe each of the four models considered herein. Table 1 defines this *c* and other variables used in the article.

**TABLE I.**
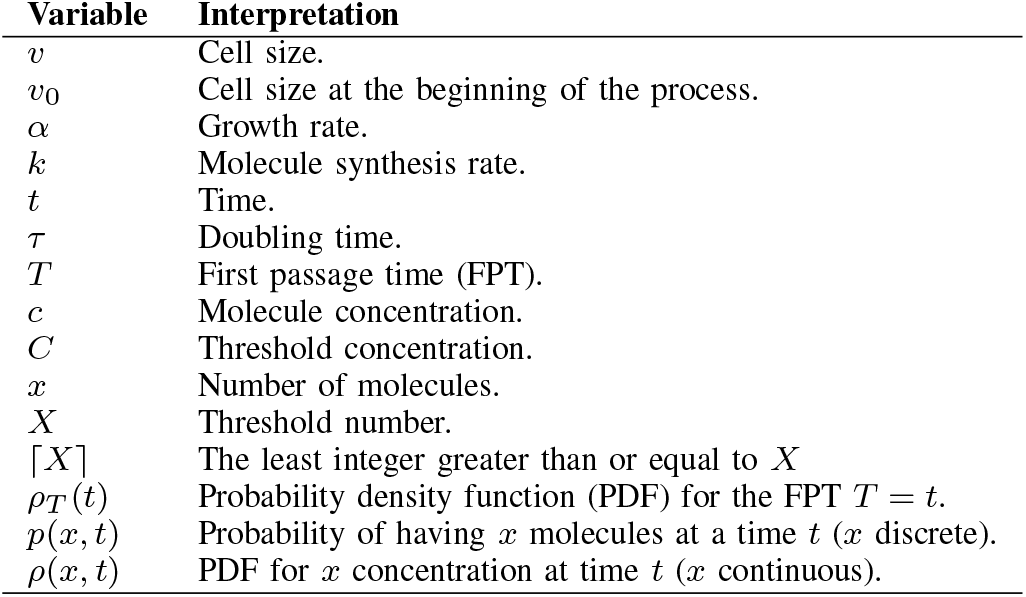
Variables employed throughout the article.

| Variable | Interpretation |
| --- | --- |
| $v$ | Cell size. |
| $v_0$ | Cell size at the beginning of the process. |
| $\alpha$ | Growth rate. |
| $k$ | Molecule synthesis rate. |
| $t$ | Time. |
| $\tau$ | Doubling time. |
| $T$ | First passage time (FPT). |
| $c$ | Molecule concentration. |
| $C$ | Threshold concentration. |
| $x$ | Number of molecules. |
| $X$ | Threshold number. |
| $\lceil X \rceil$ | The least integer greater than or equal to $X$ |
| $\rho_T(t)$ | Probability density function (PDF) for the FPT $T = t$. |
| $p(x, t)$ | Probability of having $x$ molecules at a time $t$ ($x$ discrete). |
| $\rho(x, t)$ | PDF for $x$ concentration at time $t$ ($x$ continuous). |

### A. Cell growth and gene expression

Let *v*(*t*) be the cell size at time *t*. We assume exponential growth [35], [36] described by:

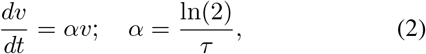

where *α* is the growth rate related to the doubling time *τ*. By solving (2), we know that size follows an exponential function of time *v*(*t*) = *v*_0_*e*^*αt*^ with *v*_0_ := *v*|_*t*=0_.

Now, consider that cell has *x* molecules. The concentration *c*, in this case is related to *x* through the size *v* by:

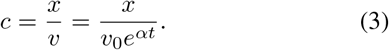

As suggested in some previous studies [20], [27], [28], we can consider the molecule number to increase at a size-dependent rate *r*(*t*) = *kv*(*t*), where *k* is a constant. In this case, the mean molecule number ⟨*x*⟩ follows [27]:

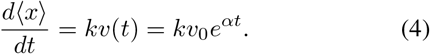

Since cells are growing exponentially, the mean concentration 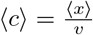, using (2) and (4), obeys:

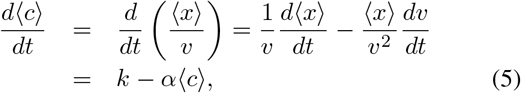

which, with the initial condition ⟨*c*⟩|_*t*=0_ = 0, it has solution

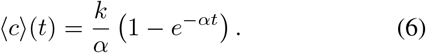

In this article, we will present some approaches to describe the random fluctuations of the process associated with (5), and how the FPT statistics change depending on each description. First, we start with three common models that do not consider explicitly the growth and a fourth model where cells grow along the time.

### B. Birth-Death Poisson Discrete Process

In a Birth-Death model, protein synthesis is described as a Poisson process with size-independent production rate *k* while degradation, dilution, and effects of size division are taken as a death process with occurrence rate *αc* (Fig. 1A). Since this model considers the size constant, the concentration *c* is equivalent to the molecule number *x*. In the same way, threshold amount *X* is identical to the threshold concentration *C*.

**Fig. 1.**
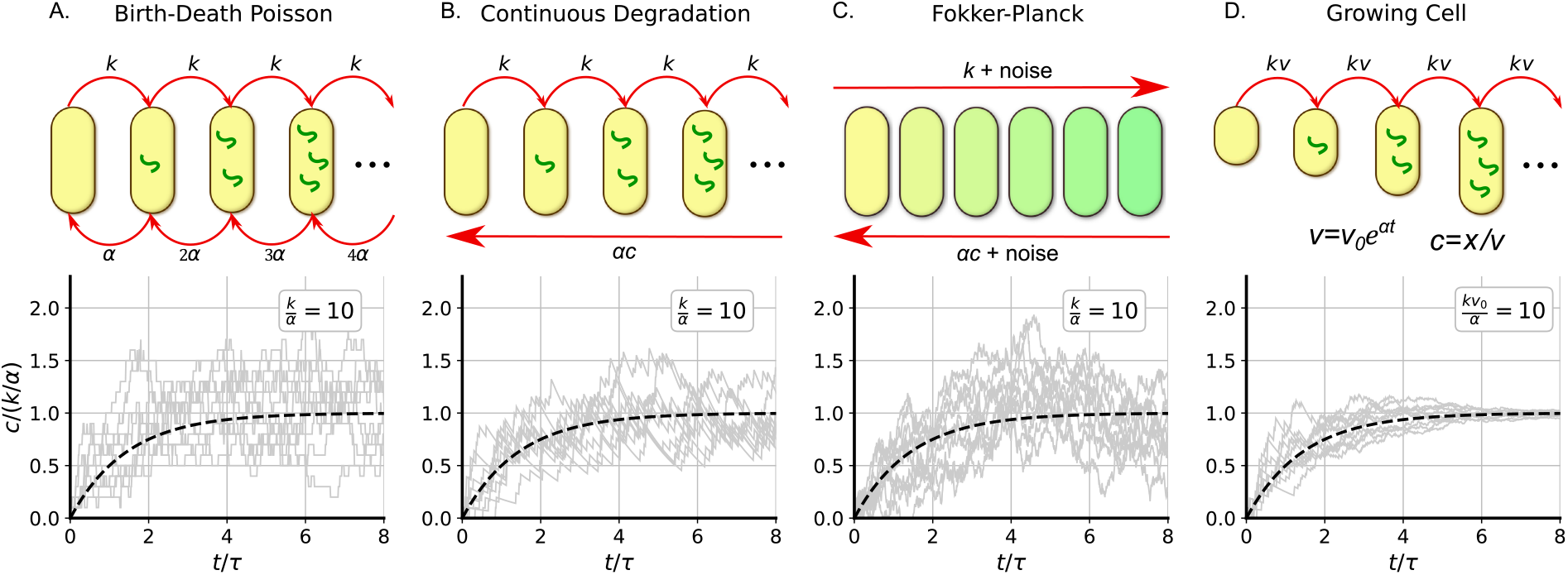
Diagram of the considered models for describing the protein accumulation. A. *Top:* Discrete synthesis of molecules inside the cells can be approximated as a Birth-Death Poisson process with the birth rate *k* and death rate *α* times the molecule number. *Bottom:* An example of some stochastic trajectories in gray. B. *Top*: Discrete synthesis at a rate *k* plus deterministic dilution at a rate *α* times the concentration *c. Bottom:* An example of some stochastic trajectories in gray.. C. *Top*: Concentration dynamics as a continuous stochastic process following the Fokker-Planck equation with production rate *k* and degradation-dilution rate *α. Bottom:* An example of some stochastic trajectories in gray. D. *Top*: Discrete protein synthesis in an exponentially growing cell. The synthesis rate is proportional to the cell size. The concentration *c* is the molecule number *x* over the size *v. Bottom:* An example of some stochastic trajectories in gray. A Black dashed line is presented in the bottom plots corresponding to the deterministic trajectory 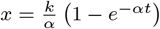, derived in (6).

To solve the FPT distribution, we set the threshold concentration *C*, let *⌈C⌉* being the least integer greater than or equal to *C*. The concentration *c* is in the set *c* ∈ *{*0, 1, *· · · ⌈C⌉}* and, considering the state *⌈C⌉* as an absorbent one:

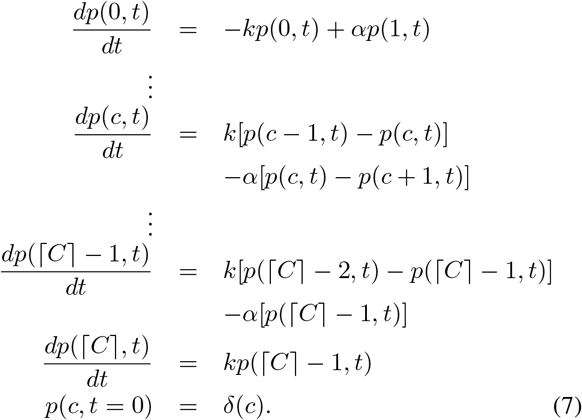

The FPT distribution *ρ*_*T*_ (*t*) can be estimated after integration of the system (7) through:

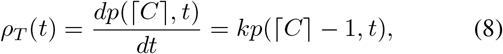

which can be used to find the *n*-th moment ⟨*T*^*n*^⟩ of *ρ*_*T*_ (*t*):

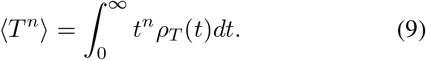

Some studies [16] obtained analytic expressions for *T*^*n*^ solving (7). In this article we are interested on study the average FPT ⟨*T* ⟩ and the variability of these FPTs measured by the squared coefficient of variation *CV* ^2^(*T*) as follows:

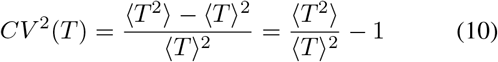

The mean FPT ⟨*T* ⟩, for example, can be compared to the FPT of a deterministic process *T*_*C*_ derived from (6):

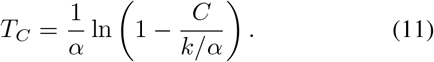

As this equation suggest, we can compare different relative dynamics by normalizing the concentration threshold *C* by *k/α*. Fig. 2A shows the dependence of ⟨*T* ⟩ for different values of 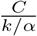. This *T*_*C*_ also defines an upper boundary for ⟨*T* ⟩ being its limit as *k/α*→ ∞ (Fig. 2A dashed line).

**Fig. 2.**
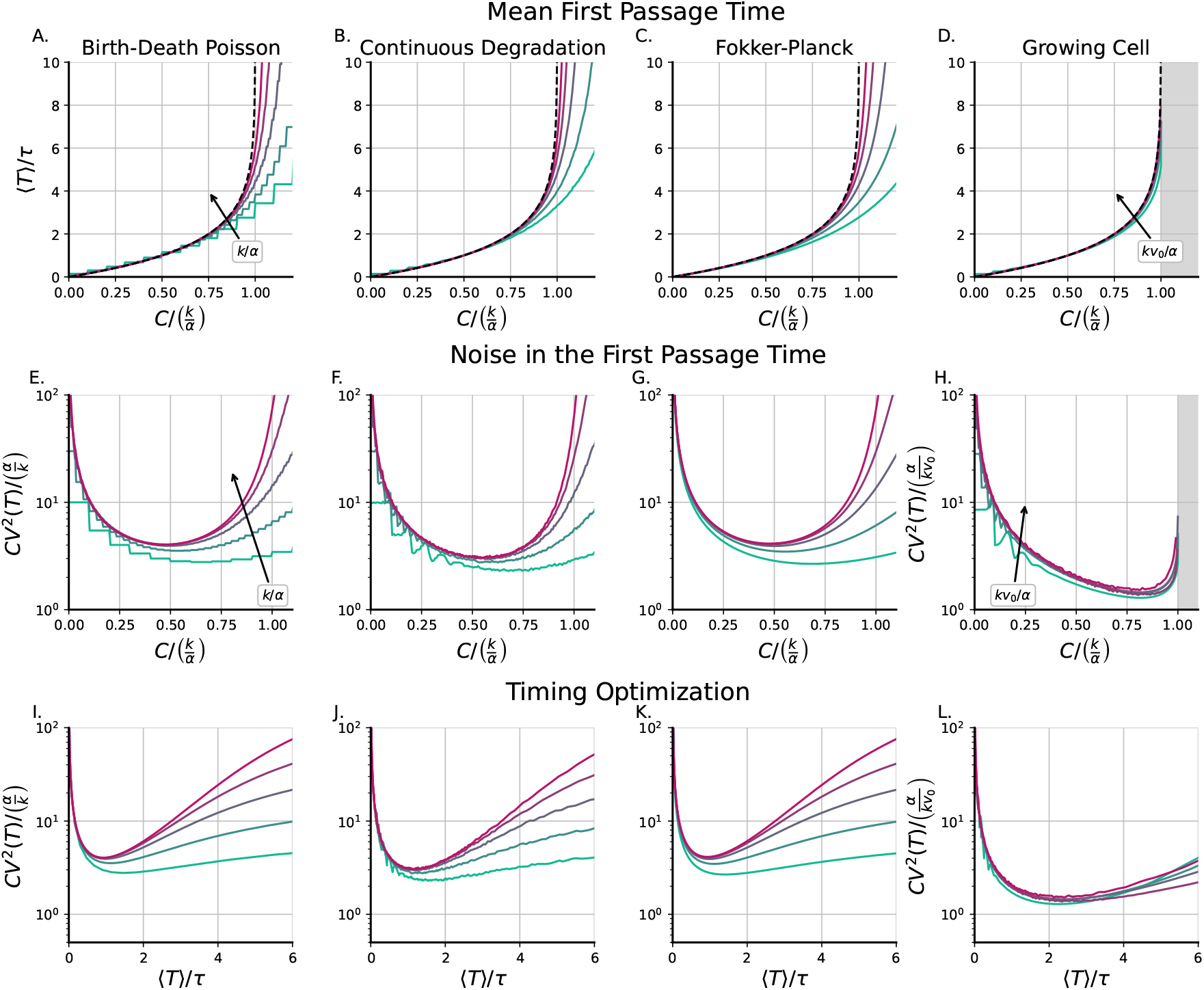
FPT statistics for different threshold concentrations considering the four models for protein accumulation. *Top*: Mean first passage time (FPT) as a function of different concentration thresholds (*C*) for the four models explored here and different noise regimes (*k/α* ∈ {10, 30, 100, 300, 1000} cyan line being *k/α* = 10 or *kv*_0_*/α* ∈ {10, 30, 100, 300, 1000} for growing cell, cyan line being *kv*_0_*/α* = 10). *Middle*: Stochastic fluctuations of the FPT measured as the squared coefficient of variation relative to the protein concentration in steady level *α/k* as a function of different concentration thresholds for the four explored models and different noise regimes. *Bottom*: Stochastic fluctuation of the FPT’s *CV* ^2^(*T*) as a function of the mean FPT with *C* as the parameter for the four models explored here and different regimes of noise.

This variable *k/α* is the mean number of molecules in steady state (*t >> α*^−1^) as (6) suggest. This variable is also quantifies the protein fluctuations. In steady conditions 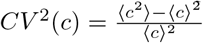 obeys [4]:

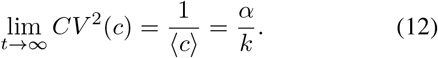

We expect that *CV* ^2^(*T*) has a similar order of magnitude as *CV* ^2^(*c*). Thus, in this article, we will present the noise in *T* in terms of *CV* ^2^(*T*)*/*(*α/k*).

### C. Discrete production and Continuous degradation

Although molecule synthesis exhibits discreteness in production, dilution by growth is a continuous deterministic process since the cell is elongating exponentially. We can model dilution by considering *c* as a continuous variable and taking the dilution as a drift term, at a growth rate *α* in the associated forward differential Chapman-Kolmogorov equation (Fig. 1B) [37]:

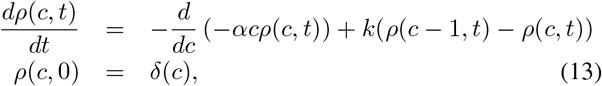

where the jumps from *c* to *c* + 1 reflect the discrete protein synthesis. Fig. 2B and Fig. 2E present results of ⟨*T*⟩ and *CV* ^2^(*T*), respectively. To estimate the solution of (13), we used the stochastic simulation algorithm as in Algorithm 1.

The result of algorithm 1 is an array of the FPT *T*_*m*_ with *m* ∈ *{*1, 2, *· · ·, M }* with *M* being the total number of simulated cells (we took *M* = 20000). From this array, we can estimate the non-central moment ⟨*T*^*n*^⟩ as:

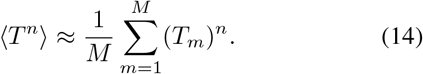

#### Algorithm 1: Stochastic Simulation Algorithm for estimating the FPT *T*_*m*_ for each of the *M* cells

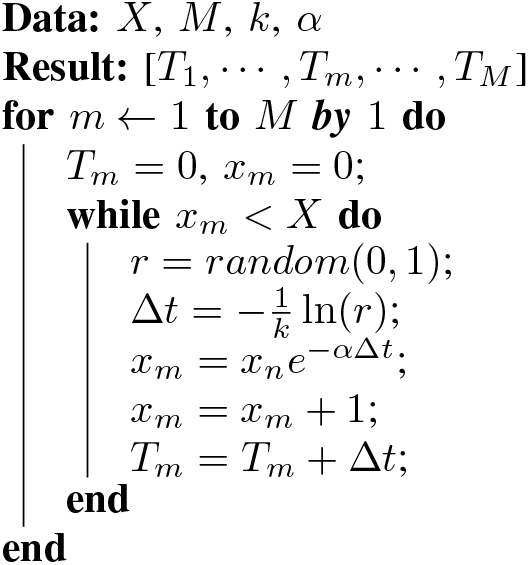

### D. Approximation of continuous stochastic dynamics: The Fokker-Planck equation

A Wiener process can approximate discrete jumps, under the assumption that the burst sizes are small relative to the molecule number (Fig. 1C) [4]. Under this approximation, *c* takes continuous values with stochastic dynamics that can be described by the SDE

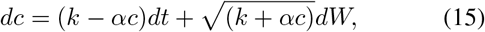

with *dW* being a random Wiener increment satisfying ⟨*dW* (*t*)*dW* (*t*^*′*^)⟩ = *δ*(*t* − *t*^*′*^) [32]. From this SDE, the distribution *ρ*(*c, t*) follows the Fokker-Planck equation

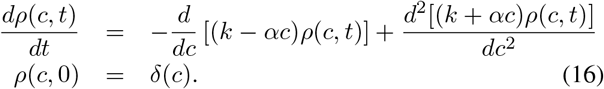

Analytic approaches for obtaining the FPT moments of *ρ*_*T*_ (*t*) are elaborated in [32] for models similar to (16). To solve numerically these moments we follow the method elaborated in Ref. [38], obtaining the moments ⟨*T*^*n*^⟩:

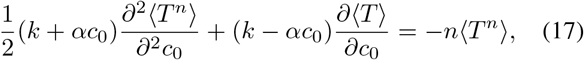

where *c*_0_ := *c*|_*t*=0_, is the concentration at the beginning of the process. We solve the problem with boundary conditions 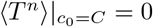 and 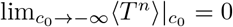 and evaluate the solution ⟨*T*^*n*^⟩ at *c*_0_ = 0. Fig. 2C and Fig. 2F show ⟨*T*⟩ and *CV* ^2^(*T*) respectively as function of *C/*(*k/α*).

### E. Size dependent propensity and continuous growth: FPT for a threshold concentration in a growing cell

This model considers that concentration *c* and molecule number *x* are related through the size *v* as shown in (3). The protein accumulation can be described using a master equation similar to (7) If *p*_*i*_(*t*) := *p*(*x* = *i, t*) is the probability for the cell having *i* molecules at a time *t*, protein accumulation is described by:

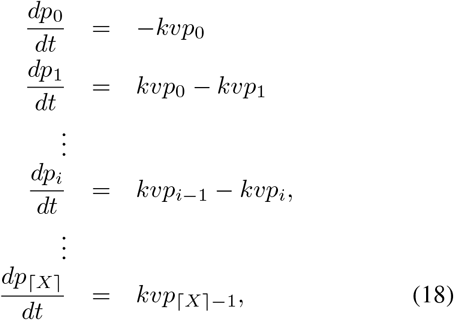

with *v* = *v*_0_*e*^*αt*^ and *⌈X⌉* being the lowest integer greater than or equal to the threshold amount *X* = *Cv*. Here, the number of molecules *x* is incremented by 1 at a rate proportional to cell size *kv*(*t*) as explained in (4).

Since *C* is fixed and cell size *v* is growing, *X* is a function of time:

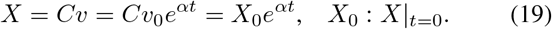

To explain the effects of having a growing *X* as in (19), consider the case where the initial volume *v*_0_ is small enough, such as *X*_0_ = *Cv*_0_ *<* 1. We can define the time *t*_1_ *>* 0 such as the instant when *X* := 1:

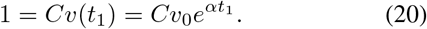

This means:

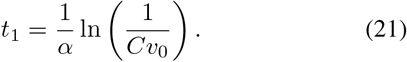

During 0 *< t < t*_1_ the system follows the master equation:

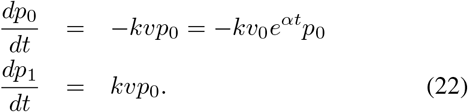

During 0 *< t < t*_1_, cells in the state *x* = 1 have already crossed the threshold. The analytic solution of (22) for *p*_1_(*t*) is given by:

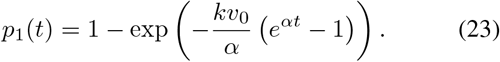

This helps us to estimate the FPT distribution:

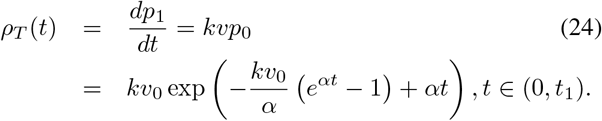

When *t*_1_ *< t < t*_2_, the remaining cells, those ones still in *x* = 0 have to jump to state *x* = *⌈X⌉* = 2 to cross the threshold concentration. The master equation is now

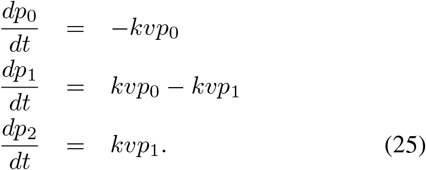

The initial conditions of the distribution *p*_*i*_ at time *t*_1_ are:

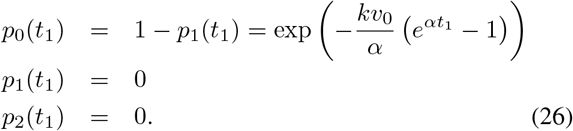

Hence, the distribution of FPTs during *t* ∈ (*t*_1_, *t*_2_) is 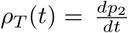 which is solved using (25).

In general, we define the time *t*_*X*_ where the cell grows so large that the threshold number is *X* molecules:

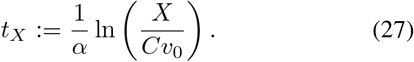

During *t*_*⌈X⌉*−1_ *< t < t*_*⌈X⌉*_, the master equation is (18) and the initial conditions, similar to (26), correspond to the distribution vector *p*_*i*_(*t*_*⌈X⌉*−1_)∀*i*.

The vector *p*_*i*_(*t*), with *t* ∈ (*t*_*⌈X⌉*−1_, *t*_*⌈X⌉*_)) and *X* = *X*_0_*e*^*αt*^, can be calculated knowing the values of *p*_*i*_(*t*_*⌈X⌉*−1_), 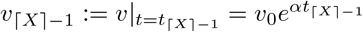, and using the recursive formula:

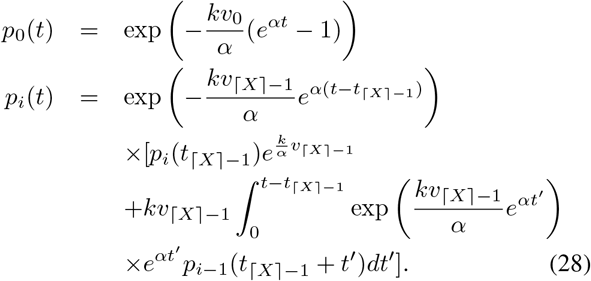

with the additional initial condition *p*_*⌈X⌉*−1_(*t*_*⌈X⌉*−1_) = 0.

After solving *p*_*⌈X⌉*_ using (28), the FPT distribution is:

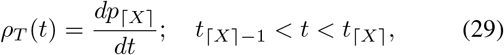

In Fig. 2D and Fig. 2H, we present numerical computations for ⟨*T*⟩ as a function of the threshold concentration *C/*(*k/α*) for different values of 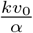. This parameter can be associated to the parameter *k/α* in the classical models. The interpretation of this ratio will be discussed in the next section.

## III. Discussion

In this work, we revisited three models of molecule accumulation by cells up to a threshold concentration. These models do not account for cell size dynamics and they consider that concentration is equivalent to molecule number. The parameter *k/α* defines the mean concentration at steady state. As explained in (12) and articles such as [4], [5], this number also defines the relative noise of the process. In the growing cell model, the regime of noise is defined by the parameter 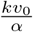. This quantity is interpreted as the number of molecules needed to reach steady concentration *k/α* when the cell has size *v*_0_. Observe that for a fixed concentration *k/α*, by changing *v*_0_, the noise of the process also changes.

In the growing cell model (Fig. 2D and 2H), we observe that the FPT moments diverge as the threshold concentration exceeds the steady value *k/α*. To understand this effect, we derived the Theorem 1 in the appendix and present some stochastic trajectories as examples (Fig. 1D). As the theorem 1 explains, as time passes, the concentration fluctuations decrease exponentially. As a result, some cells can cross thresholds higher than the steady concentration at the beginning of the process. However, cells that did not cross this threshold have less probability of doing it in the future.

In previous studies [16], [32], protein accumulation FPT problem focused on *E. coli* lysis by lambda phage. Once the process starts, the protein holin is accumulated on the bacterial membrane and punctures it after its concentration crosses a threshold concentration [39]. Some evolutionary studies observed that the parameters of the chemical process evolved such as the lysis occurs at a concentrations with an associated FPT having minimal fluctuations [40], [41].

Considering this experimental context, we can show (Fig. 2I, 2J, 2K and 2L) the profile *CV* ^2^(*T*) vs ⟨*T*⟩ predicted by each model. In classical models, *CV* ^2^(*T*) shows a a global minimum close to ⟨*T*⟩ ≈ *τ* while in the model of growing cell, this minimum corresponds to a larger FPT ⟨*T*⟩ ≈ 2*τ*. As discussed above, the fact that in growing cells the fluctuations are progressively smaller can also explain why the position of the optimum noise time is higher in the growing cell model than in the classical ones.

## IV. Conclusions

This article explores the statistics of the first passage times (FPT’s) for the process of protein accumulation up to a threshold concentration. We revisited three classical models as a context: the Birth-Death Poisson Process, the discrete production with continuous degradation, and the Fokker-Planck approximation. We compare their results on FPT statistics with a fourth model proposed by us, which considers a growing cell with protein produced at a rate proportional to the cell size. We see how the first three models are equivalent in these timing statistics and how they differ from the last model.

The first three models predict that, on average, cells can reach concentrations higher than the mean concentration at steady-state. On the other hand, the growing cell model predicts that all the FPT moments diverge if the threshold concentration is higher than this steady concentration.

Regarding the timing variability, we observe how the growing cell model predicts fewer fluctuations in timing than the classical models. After finding the average FPT that minimizes these timing fluctuations, this first model predicts an optimum FPT almost twice the one found by the other models.

## ACKNOWLEDGMENT

AS is supported by NIH 1R01GM124446-01.

## APPENDIX

### Theorem 1

If the dynamics of the vector 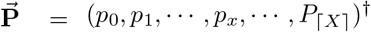 is described by the system:

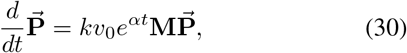

with *k, v*_0_ and *α* constants, and **M** a matrix of constant elements *M*_*i,j*_ with properties:

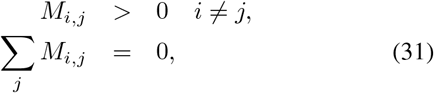

and defining the concentration *c* := *x/*(*v*_0_*e*^*αt*^), then, the fluctuations in *c* decay asymptotically as an exponential function of time, this is:

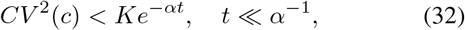

for a constant *K*.

*Proof:* Consider the mapping:

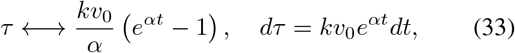

which after been applied to (30), reduces the system into a master equation of a Poisson process:

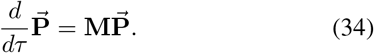

which has the property:

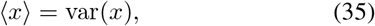

for any *τ*, with ⟨*x*⟩ = Σ _*x*_ *xp*_*x*_ and var(*x*) = Σ _*x*_ *x*^2^*p*_*x*_ − (Σ_*x*_ *xp*_*x*_)^2^. Since the mapping (33) is monotonically increasing function of the time and state independent, this property also holds for the system (30).

Given that the protein *x* is produced at a rate proportional to the size, the mean protein follows:

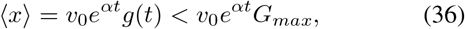

with *G*_*max*_ a constant since *g*(*t*) is bounded up to time *t*. Now, using (35), we obtain:

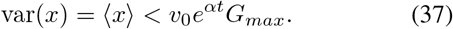

While the mean concentration follows

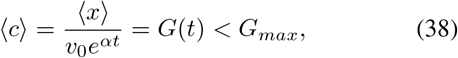

the fluctuations of *c* = *k/*(*v*_0_*e*^*αt*^) satify:

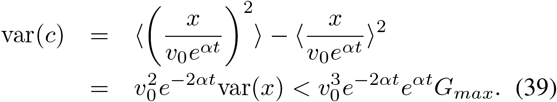

Now, the squared coefficient of variation follows:

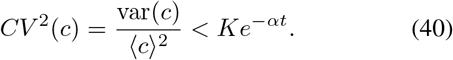

▪

## Notes

### Competing Interest Statement

The authors have declared no competing interest.

